# Linear versus deep learning methods for noisy speech separation for EEG-informed attention decoding

**DOI:** 10.1101/2020.01.22.915181

**Authors:** Neetha Das, Jeroen Zegers, Hugo Van hamme, Tom Francart, Alexander Bertrand

## Abstract

**Objective:** A hearing aid’s noise reduction algorithm cannot infer to which speaker the user intends to listen to. Auditory attention decoding (AAD) algorithms allow to infer this information from neural signals, which leads to the concept of neuro-steered hearing aids. We aim to evaluate and demonstrate the feasibility of AAD-supported speech enhancement in challenging noisy conditions based on electroencephalography (EEG) recordings.

**Approach:** The AAD performance with a linear versus a deep neural network (DNN) based speaker separation was evaluated for same-gender speaker mixtures using 3 different speaker positions and 3 different noise conditions.

**Main results:** AAD results based on the linear approach were found to be at least on par and sometimes even better than pure DNN-based approaches in terms of AAD accuracy in all tested conditions. However, when using the DNN to support a linear data-driven beamformer, a performance improvement over the purely linear approach was obtained in the most challenging scenarios. The use of multiple microphones was also found to improve speaker separation and AAD performance over single-microphone systems.

**Significance:** Recent proof-of-concept studies in this context each focus on a different method in a different experimental setting, which makes it hard to compare them. Furthermore, they are tested in highly idealized experimental conditions, which are still far from a realistic hearing aid setting. This work provides a systematic comparison of a linear and non-linear neuro-steered speech enhancement model, as well as a more realistic validation in challenging conditions.

## 1. Introduction

In a noisy environment with multiple speakers talking simultaneously, i.e., the so-called cocktail-party problem, a person with normal hearing has the ability to focus attention on one speaker and ignore the other speakers and surrounding noise in a seemingly effortless manner. People with a hearing impairment, on the other hand, find such situations extremely challenging. Although modern hearing aids allow to suppress background noise, a major unsolved problem is to determine which of the speakers is the desired one, and which speakers should be treated as noise. Existing approaches use unreliable heuristics based on speaker loudness or other acoustic features or simply assume that the target speaker is always in the frontal direction.

Recent developments in the field of neuroscience have shown that it is possible to decode the auditory attention of a listener in a multi-talker environment from brain signals recorded with magneto- or electro-encephalography (M/EEG) [1–3]. This opens up new opportunities to design smarter hearing devices, augmented with electrodes to record neural signals, that decode in real time to which speaker a listener is attending, thereby assisting the hearing device to determine and enhance the attended speaker.

The envelope tracking of speech streams in a listener’s cortical responses, and more so of the attended speech stream, is well established in the literature [4–6]. Furthermore, this differential tracking of attended and unattended streams is also present in hearing impaired listeners [7–9]. This fact can be exploited to design algorithms that perform auditory attention decoding (AAD). Various methods have been developed to achieve AAD, addressing different kinds of decoders [2, 10–13], stimulus features [3, 14, 15], data acquisition [16–18], electrode selection and miniaturization strategies [16, 18, 19], etc.

The aforementioned AAD studies in [2–19] assume access to the clean speech signals to perform AAD, which are not available in a practical setting. Indeed, a hearing aid only has access to noisy microphone recordings in which multiple speech sources and noise sources are mixed. The first study that aimed for AAD without access to the clean speech streams was published in [20], where a multiplicative non-negative ICA (M-NICA) algorithm [21] was used to extract the individual speech envelopes from a noisy multi-microphone input. The AAD method in [14] was then used to select the attended speaker, which was then extracted from the microphone array using a multi-channel Wiener filter (MWF) [20]. This pipeline was later extended in [22] with a twofold MWF (one for each speaker) leading to better decoding performance and less variation across different acoustic conditions. However, while in both of these studies, the performance of the speaker separation algorithm was evaluated for a range of background noise levels and speaker positions, the EEG signals used for AAD were collected from subjects who listened to a 2-speaker scenario with 180° speaker separation and without background noise. Therefore, there was a mismatch between acoustic conditions in the AAD experiments and the audio processing modules. In [23], speech enhancement was achieved using binaural direction-of-arrival (DOA) estimators with linearly constrained minimum variance (LCMV) beamformers. The AAD module used EEG signals recorded in two relatively high signal-to-noise ratio (SNR) conditions and two reverberation conditions.

The audio processing in [20, 22, 23] is based on linear beamforming and linear source separation techniques, which do not require any training data and have the advantage of being computationally cheap. In recent years, DNN-based approaches have become a popular alternative to solve the speaker separation problem, particularly for the challenging single-microphone scenario [24–26] and the even more challenging scenario with additional background noise [27, 28]. When using multiple microphones, separation performance can be increased by making use of the spatial information of the sources [29, 30]. In [31, 32], a single-microphone DNN-based approach was used for speaker separation in an AAD context. However, both of these studies involved training and testing in noise-free conditions. Furthermore, AAD was performed on electro-corticography (ECoG) data, which is an invasive approach involving surgery. This is a limiting factor for hearing-aid users. Non-invasive techniques like EEG are more preferable in this regard.

It is evident that a detailed analysis of neuro-steered speech enhancement under challenging acoustic conditions is necessary to draw conclusions about its feasibility. Despite the promising results in the aforementioned studies, all of them have important caveats, which may lead to overoptimistic conclusions in a context of demonstrating viability of neuro-steered speech enhancement algorithms. To summarize: The audio processing of [20,22] was tested in various noisy conditions with different angles between speakers, but the EEG recordings were performed in a noise-free condition with 180° separation, resulting in a mismatch between the acoustic conditions of the microphone and neural signals. This mismatch does not occur in [23], but only mild noisy conditions are considered and the speaker positions are the same in all conditions. Finally, it is noticed that in [32] ECoG is used instead of EEG. Furthermore, the authors only report results in noiseless conditions, using single-microphone recordings(while most state-of-the-art hearing aids contain multiple microphones), and with competing speakers of different gender‡.

In addition to these practical limitations, the different settings across the aforementioned studies make it impossible to draw conclusions about the impact on AAD performance of non-linear (DNN-based) methods like [31, 32], and linear methods like [20, 22, 23], as well as the effect of using spatial information from multiple microphones versus single-microphone methods.

The contribution of this paper is twofold. First, the aforementioned caveats are avoided by focusing on realistic scenarios involving microphone recordings from a binaural hearing aid in challenging noisy conditions with same-gender competing talkers at various relative speaker positions. In each acoustic condition, the audio and neural data are matched to each other, i.e., the audio signals presented to the subjects during the EEG recording are the same as those used for the audio signal processing. Secondly, the potential improvement of using a DNN-based speech separation algorithm compared to a computationally cheap and training-free linear signal processing algorithm is investigated. The performance of the extended linear approach is shown to be at par or better than the purely DNN-based approach. The potential advantage of using a combination of both approaches, i.e. a DNN-based speech separation to support a linear beamformer, is also investigated. The superior performance of such a system that combines *the best of both worlds* is demonstrated in terms of improvement in signal-to-interference-plus-noise ratio (SINR), perceptual evaluation of speech quality (PESQ) [33] and AAD accuracy. The study focuses on two representative state-of-the-art algorithms, i.e., deep clustering [24] as the DNN-based approach and M-NICA+MWF [22] as the linear method. In both cases, a multi-microphone input is used, which will substantially improve speech separation compared to a single-microphone input.

The outline of this paper is as follows. The different neuro-steered speech enhancement pipelines are presented in Section 2. The various experiments used to validate the different approaches are presented in Section 3. In Section 4, the results of the experiments are shown, and the various implications of these findings are discussed in Section 5.

## 2. Methods

The proposed neuro-steered speech enhancement pipeline is shown in the block diagram of Figure 1 for a 2-speaker scenario. Although the analysis in this paper focuses on a 2-speaker scenario, the speech enhancement procedure in Figure 1 can straightforwardly be generalized to a larger number of speakers (denoted by *N* in the text). The pipeline consists of three steps:

**Figure 1:**
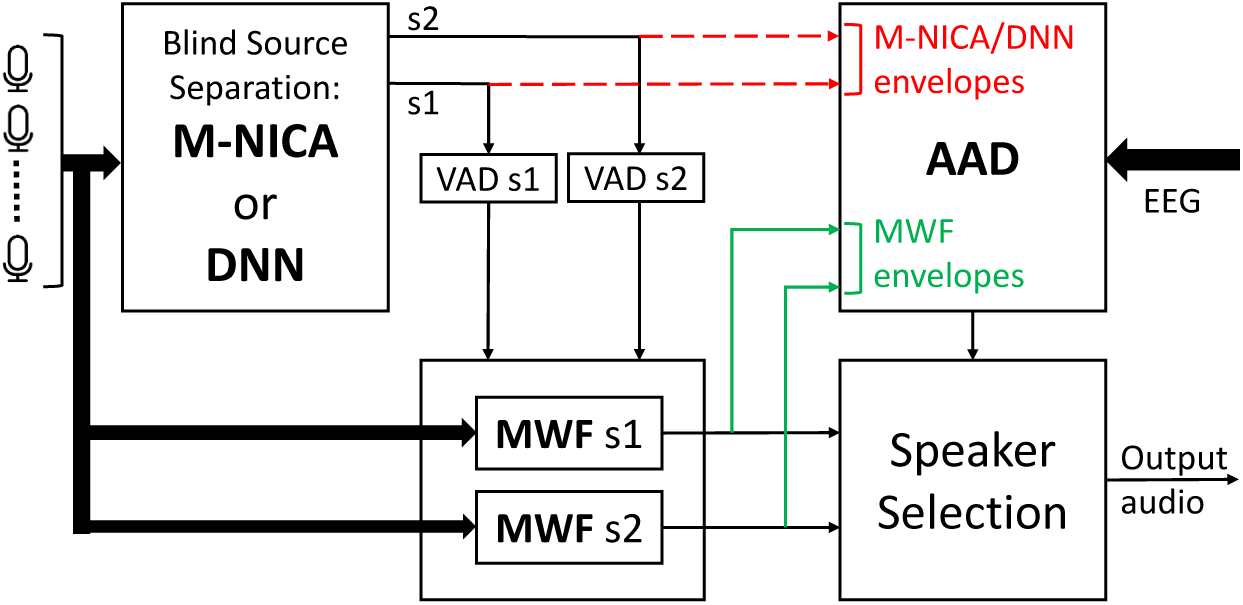
Block diagram of the proposed neuro-steered speech enhancement algorithm for a 2-speaker scenario, consisting of modules for blind source separation (BSS), filtering for speech enhancement (using MWF) and auditory attention decoding. The red (dashed) track and the green track correspond to two different implementations, as explained in the text.

i. In the M-NICA/DNN module, the speech sources are separated in a fully ‘blind’ (unsupervised) fashion. Optionally, the estimated speech envelopes obtained from the M-NICA/DNN module can be used as side information for the semi-supervised MWF module, which typically achieves a better separation and denoising performance. This will be discussed in Subsection 2.2.
ii. In the AAD module, the attended speech envelope from the listener’s EEG is reconstructed using a pre-trained decoder. This reconstructed speech envelope is correlated with the speech envelopes of the individual speakers extracted in step 1, where the speaker with the highest correlation is selected as the attended speaker. To this end, either the envelopes from the M-NICA or DNN algorithm can be used directly (red dashed lines in Figure 1), or a new set of envelopes can be extracted from the denoised MWF outputs (green lines in Figure 1). Both cases will be investigated, where the envelopes in the red dashed lines will be referred to as M-NICA or DNN envelopes (depending on which algorithm is used in the BSS block) and the ones in the green lines as MWF envelopes. More details about the AAD are given in Subsection 2.1.
iii. Once the speaker has been decoded, the corresponding MWF output signal is selected as the final output, which ideally consists of a denoised version of the speech signal corresponding to the attended speaker.

In the remainder of this paper, we assume that the number of speakers *N* is known. In practice this would require a supporting algorithm to estimate *N* from the microphone recordings, e.g., based on subspace analysis, which is beyond the scope of this paper.

### 2.1. Auditory attention decoding (AAD)

In order to choose which enhanced speech estimate will be presented as output of the neuro-steered speech enhancement pipeline, it is necessary to know which speech signal is being attended to by the listener. Towards this goal, an AAD module is used which consists of a pre-trained spatio-temporal decoder (subject-specific training) which takes the listener’s EEG data as an input to reconstruct the speech envelope of the attended speaker. In the training data, the ground truth about the subject’s attention is known, and the decoder is designed such that the difference between its output and the attended speech envelope is minimized in the mean square error (MSE) sense. The algorithm described in [14] was used for our decoder design. The reconstructed attended speech envelope is given by

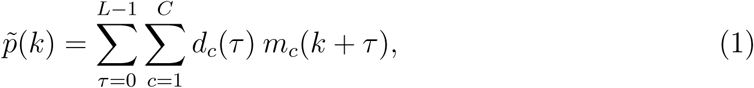

where *m*_*c*_(*k*) is the value of the *c*-th EEG channel at sample time *k*, and *d*_*c*_(*τ*) denotes the decoder weight at the *c*-th channel at time lag *τ*. All decoder weights are stacked in a vector ***d*** = [*d*_1_(0), *d*_1_(1), …, *d*_1_(*L* − 1), *d*_2_(0), …, *d*_2_(*L* − 1), …, *d*_*C*_(0), …, *d*_*C*_(*L* − 1)]^*T*^. Stacking EEG samples of all *C* channels for *L* time lags, we get ***m***(*k*) = [*m*_1_(*k*), *m*_1_(*k* +1), …, *m*_1_(*k* + *L* − 1), *m*_2_(*k*), …, *m*_2_(*k* + *L* − 1), …, *m*_*C*_(*k*), …, *m*_*C*_(*k* + *L* − 1)]^*T*^ ∈ ℝ^*LC*^.

Using this notation, (1) can be written as 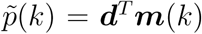. The optimal decoder that minimizes the MSE between the reconstructed attended envelope 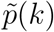 and the attended speech envelope *p*_*a*_(*k*) is given by

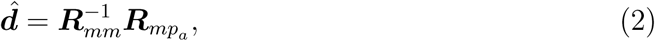

where ***R***_*mm*_ = *E*{***m***(*k*)***m***(*k*)^*T*^} ∈ ℝ^*LC*×*LC*^ is the EEG covariance matrix, 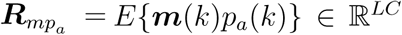 is the cross-correlation vector between the EEG data and the attended speech envelope, which is known during the training of the decoder and *E*{.} denotes the expected value operator. At test time, the correlation coefficients between 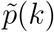 and the estimated speech envelope of each speaker are then computed. The speech envelope that results in the higher correlation is selected as the attended speech stream.

### 2.2. Blind source separation and denoising

The microphone signal of a mixture of *N* speakers can be represented in the short-time Fourier transform (STFT) domain as

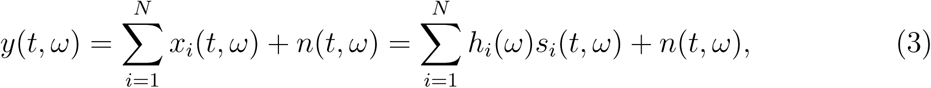

where *t* indicates the time, *ω* the frequency index, *n*(*t, ω*) represents the noise recorded by the microphone and *x*_*i*_(*t, ω*) represents the observed signal of the *i*-th speaker *s*_*i*_, as recorded by the microphone. The transfer function that models the acoustic propagation path between the source *s*_*i*_ and the microphone is given by *h*_*i*_(*ω*).

In a multi-microphone setting, such as a (binaural) hearing device, where *M* microphones are present, (3) generalizes to

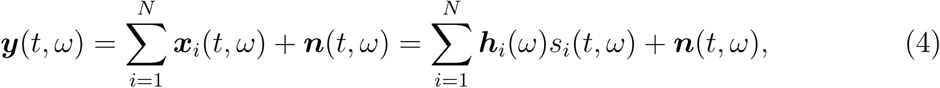

where ***y***(*t, ω*) = [*y*_1_(*t, ω*), *y*_2_(*t, ω*), …, *y*_*M*_ (*t, ω*)]^*T*^ represents the microphone components stacked over *M* microphone channels, ***h***_*i*_(*ω*) = [*h*_*i*,1_(*ω*), *h*_*i*,2_(*ω*), …, *h*_*i,M*_ (*ω*)]^*T*^ with *h*_*i,j*_(*ω*) modeling the path between the source *s*_*i*_ and microphone *j*, and ***n***(*t, ω*) = [*n*_1_(*t, ω*), *n*_2_(*t, ω*), …, *n*_*M*_ (*t, ω*)]^*T*^ represents the stacked components of noise in the *M* microphones. A multi-microphone mask (or a beamformer) ***w***_*i*_(*t, ω*) can be estimated such that the speech signal can then be estimated as

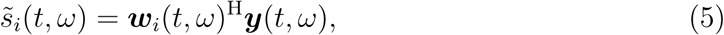

where superscript H denotes the conjugate transpose operator. Note that, while the symbol 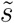 is used in (5) for the sake of clarity, the goal is not necessarily to perform a dereverberation, i.e., 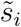 may be closer to the recorded speech signal *x*_*i*_(*t, ω*) = *h*_*i*_(*ω*) * *s*_*i*_(*t, ω*) than to the ‘dry’ speech signal *s*_*i*_ itself.

Two methods are presented to estimate the mask in (5): one that is fully linear, based on the M-NICA algorithm, and one that makes use of a DNN.

i. The linear multi-channel signal processing approach using M-NICA provides an estimate of the energy envelope of each individual speaker (see Subsection 2.2.1). These envelopes can be directly fed to the AAD module (red dashed track in Figure 1 as in [34], or alternatively, they can be used as a voice activity detection (VAD) mechanism to design an *N* -fold MWF which estimates per-speaker masks ***w***_*i*_ in (5), as explained in Subsection 2.2.2. The envelope of 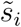 from (5) can then be fed to the AAD module as in [22] (green track in Figure 1).
ii. On the other hand, a DNN can be trained to separate speech mixtures into individual speech streams (see Subsection 2.2.3). As opposed to M-NICA, the DNN directly estimates the masks ***w***_*i*_ to compute (5), after which the speech envelopes can be extracted and directly fed to the AAD module as in [31, 32] (red dashed track in Figure 1). Alternatively, the envelopes obtained from the DNN can also be used as a VAD to design MWF masks (green track in Figure 1).

#### 2.2.1. Multiplicative non-negative independent component analysis (M-NICA)

The M-NICA algorithm was introduced in [21], and has been used to blindly separate speech envelopes from a multi-microphone input [20, 34]. The M-NICA algorithm operates in the short-term energy domain, thereby transforming the problem into a (non-negative) energy envelope separation problem. To this end, the short-term energy signal from each microphone signal is computed over a window of *K* samples. The M-NICA is applied on the resulting *M* energy signals (note that M-NICA operates at a sampling rate that is *K* times lower than the microphone signal sampling rate). For more details on the implementation of M-NICA for demixing speech envelopes, we refer to [21, 34].

M-NICA exploits the inter-channel differences in the energy distributions of the speech sources in order to extract the energy envelopes of each individual speaker. In the case of a binaural hearing aid, these differences are mostly due to head shadow effects. The non-negativity of the underlying sources facilitates the use of only second order statistics in its computations, in comparison to other source separation algorithms that use higher order statistics. This, together with the *K*-fold reduction in sampling rate, leads to a low computational complexity which is a desirable factor to incorporate such a source separation scheme in an actual hearing prosthesis.

#### 2.2.2. MWFs for source separation and/or denoising

In this paper, an *N* -fold MWF (one for each speaker) will be used to estimate each speech signal in real time. The MWF is a data-driven and adaptive linear beamforming algorithm which allows to efficiently denoise and extract a single speaker from a mixture by computing the mask in (5) which results in the linear estimate with the smallest MSE, i.e., the linear minimum MSE (LMSSE) estimate. It is a semi-supervised method as it requires a voice activity detector (VAD) for the target speaker, i.e., it needs to know at which times the target speaker is not active in order to estimate the second-order statistics of the background noise and interfering speakers. The VAD information can be straightforwardly extracted by thresholding the speech envelopes obtained from the M-NICA algorithm (or alternatively using the deep learning algorithm as explained in Subsection 2.2.3). The *i*-th MWF computes the LMMSE mask ***w***_*i*_(*t, ω*) in (5) that yields the estimate 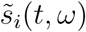 that is closest to 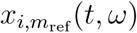, i.e., the *i*-th speaker’s contribution in an arbitrarily-chosen reference microphone *m*_ref_. The MWF ***w***_*i*_(*t, ω*) is defined as

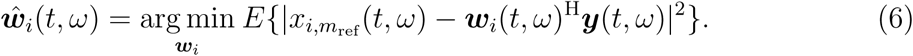

The solution for this LMMSE problem is given by [35]:

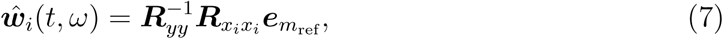

where ***R***_*yy*_ = *E*{***y***(*t, ω*)***y***(*t, ω*)^H^}, 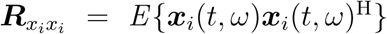, and 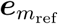 denotes the *m*_ref_-th column of an *M* × *M* identity matrix, which selects the column of 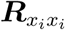 corresponding to the reference microphone. The ‘speech plus interference’ autocorrelation matrix ***R***_*yy*_ can be estimated during the time periods when the *i*-th speaker is active§. Assuming independence between all sources, 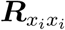 can be estimated as 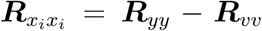. Here, ***R***_*vv*_ = *E*{***n***(*t, ω*)***n***(*t, ω*)^H^} + Σ_*j* ≠*i*_ *P*_*j*_(*ω*)***h***_*j*_(*ω*)***h***_*j*_(*ω*)^H^ is the interference autocorrelation matrix, where ***P***_*j*_(*ω*) = *E*{|*s*_*j*_(*t, ω*)|^2^}. ***R***_*vv*_ can be estimated by averaging over all STFT frames in which speaker *i* is not active. Estimating 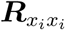 as ***R***_*yy*_ *-* ***R***_*vv*_ can result in poor filters, particularly, if ***R***_*vv*_ contains non-stationary sources. Therefore, in our computations, 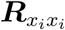 was estimated based on a generalized eigenvalue decomposition (GEVD) of ***R***_*yy*_ and ***R***_*vv*_ (details in [36]) to ensure robustness. Note that, while speech is known to be highly non-stationary, the expected values here are calculated over a long-term window, thereby capturing the average long-term power spectrum of the speech. Therefore, the MWF will not react to short-term changes in the speech spectra, it mainly focuses on the (fixed or slowly varying) spatial coherence across the microphones. To also exploit the fast spectral changes in the speech, a single-channel Wiener filter which uses short-term statistics can be added as a postfilter [37], yet this is beyond the scope of this paper.

To separately estimate ***R***_*yy*_ and ***R***_*vv*_, the MWFs require the voice activity information of the speaker it must enhance. The energy envelopes from the output of the source separation algorithm (M-NICA or a DNN approach, see subsections 2.2.1 and 2.2.3, respectively) are used to identify the active and silent periods of each speaker, for e.g., through a thresholding operation ‖, and hence form the VAD tracks that each MWF needs.

Note that the *N* MWFs work in parallel to enhance the speech streams of *N* different speakers. It is important to note that these MWFs can share a large part of the computations, as the estimation of the matrix inverse 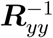 is a common requirement across them. The real-time aspects of such a system will be discussed in Subsection 5.4.

#### 2.2.3. Deep learning for source separation

Deep learning solutions for source separation differentiate between single-microphone and multi-microphone approaches as well as between methods that can and cannot cope with additional background noise. The deep learning based separation technique that will be used in this paper is called Deep Clustering (DC) [24], which can be extended to cope with noise [27, 28] and to use multi-microphone information [29].

In DC, a *D*-dimensional embedding vector ***v***(*t, ω*) is constructed for every time-frequency bin as ***v***(*t, ω*) = *f*_*tω*_(|*Y* |), where *Y* is used to refer to the complete input signal across all time-frequency bins, ***v***(*t, ω*) is a *D*-dimensional embedding vector normalized to unit length, and the function *f* is chosen to be a DNN. A (*T*Ω × *D*)-dimensional matrix ***V*** is then constructed from these embedding vectors, where *T* and Ω denote the total number of time and frequency bins, respectively. Similarly, a (*T*Ω × *N*)-dimensional target matrix ***Z*** is defined. If target speaker *i* is the dominant speaker for bin (*t, ω*), then *z*_*i*_(*t, ω*) = 1, otherwise *z*_*i*_(*t, ω*) = 0. Speaker *i* is dominant in a bin (*t, ω*) if *i* = arg max_*j*_ (|*s*_*j*_(*t, ω*)|). A permutation independent loss function is then defined as

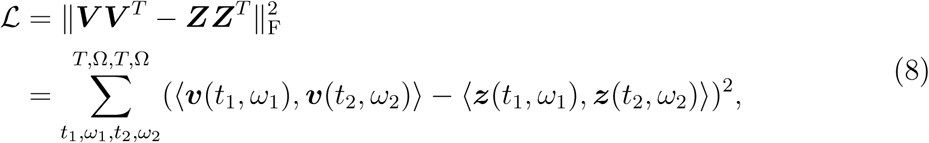

where 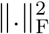 is the squared Frobenius norm. Minimizing this loss function pushes embeddings together if their time-frequency bin are dominated by the same speaker and otherwise the embeddings should be perpendicular.

After estimating ***V***, all embedding vectors are clustered into *N* clusters using k-means clustering. Speaker masks are then constructed as follows

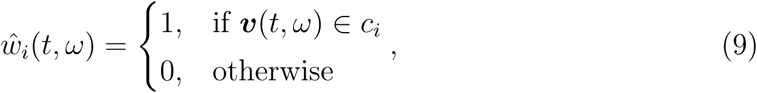

with *c*_*i*_ a cluster. These masks can be used to estimate the source signals by applying them to the original speech mixture:

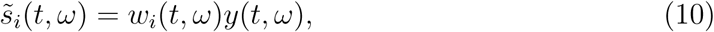

where (10) can be seen as the single channel simplification of (5). Since binary masks are estimated, it is assumed that only one speaker is active per time-frequency bin. Since speech spectra are sparse, this assumption holds to some extent [39].

Deep clustering, as presented above, is not well suited for noisy environments. The binary masks in (9) will cause the noise energy to be distributed over the speaker signal estimates. The noise energy can be filtered out by estimating an additional speech-to-noise ratio mask *α*(*t, ω*) [27]. Both the embeddings ***v***(*t, ω*) and the ratio mask *α*(*t, ω*) can be estimated by the DNN from the magnitude spectrum of the microphone signal: [***v***(*t, ω*), *α*(*t, ω*)] = *f*_*tω*_(|*Y* |). During training of the network, the target noise suppression mask is set to be

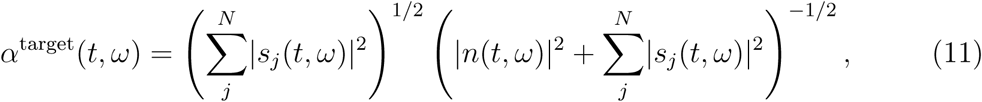

as it was shown to perform well in many noisy scenarios [40]. The loss function of (8) can then be extended to:

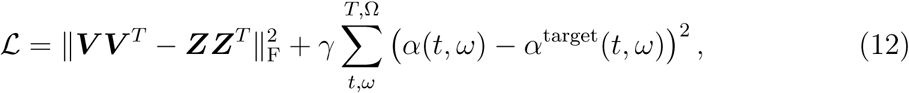

where the hyper-parameter *γ* can be tuned to weigh the importance of speech separation and noise suppression. The estimated mask of (9) is then changed to

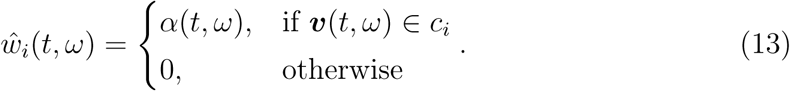

DC has been developed for single microphone setups, but can easily be extended to include spatial information ***Y***_sp_ from a multi-microphone setup [29]. This spatial information is simply appended to the input of the model to estimate the embeddings (and the noise mask): [***v***(*t, ω*), *α*(*t, ω*)] = *f*_*tω*_(|***Y*** |, ***Y***_sp_). The interchannel phase difference (IPD) between two microphones *m*1 and *m*2 is chosen to represent the spatial information [29]:

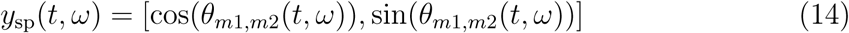

with *θ*_*m*1,*m*2_(*t, ω*) = ∠*y*_*m*1_(*t, ω*) − ∠*y*_*m*2_(*t, ω*) and ***Y***_sp_ is the matrix containing the spatial information *y*_sp_(*t, ω*) over all time-frequency bins (*t, ω*). In case there are more than two microphones (*M* > 2), the spatial features can be determined for all microphone pairs and stacked as a single feature representation.

A single mask for each speaker *i* is produced, in the same way as was done in (13) which is then copied and applied to each microphone. The mask is applied to the average of all microphone spectrograms, or equivalently¶

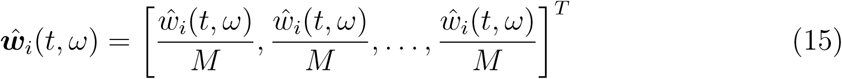

when using (5).

So far we have focused on estimating masks to filter out the interfering speaker (and noise). The estimated 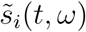 can be further enhanced in a secondary stage as in [41]. To this end, a new DNN is trained, denoted by the function *g*′, which estimates an enhanced mask for speaker *i* based on |*Y* | and the previous estimate 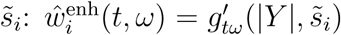. Notice that the network used for *g*′ is shared over the speakers. In the multi-microphone case, the IPD is again included to obtain 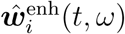 similarly to (14)-(15).

In the next step, the envelopes of the speech sources as estimated from the mask (15) can be used for auditory attention decoding (red dashed lines in Figure 1). There is also the option of using these envelopes to first compute the VADs for the *N* -fold MWFs as discussed in Subsection 2.2.2, and then using the envelopes from these MWF outputs for attention decoding instead (green lines in Figure 1). This would correspond to the algorithm proposed in [22], where the M-NICA block is replaced with a DNN. A final option, which will not be further discussed in this paper, is to use the masks estimates *ŵ*_*i*_(*t, ω*) as a weight when determining ***R***_***yy***_ and ***R***_***vv***_ in Subsection 2.2.2, similarly as was done in [42]. Initial experiments with this method gave no significant improvements in AAD accuracies.

## 3. Experiments

### 3.1. Experiments setup

#### 3.1.1. AAD setup

18 normal-hearing subjects (of which 6 males) between 20 and 25 years old participated in the experiment, where each subject had to focus on the speech stream from a particular direction, in the presence of a competing speaker from another direction and background noise. Their 64-channel EEG was recorded at a sampling rate of 8192 Hz. This dataset consists of 138 minutes of EEG recordings per subject. Further details can be found in [43]. The stimuli and the corresponding acoustic conditions used in this experiment are described below, in Subsection 3.1.2.

#### 3.1.2. RBM Dataset

The RadioBoeken Mix (RBM) dataset was built by mixing parts of 8 Dutch stories narrated by 8 different female narrators [44], taken 2 at a time as competing speech streams in the presence of different levels of background multi-talker (‘babble’) noise. The background babble noise consisted of 9 different 4-talker babble streams (2-male and 2-female) perceived to be coming from 9 equidistant positions (from -180° to 140° in steps of 40°) around the listener. The babble was constructed from 36 audiobooks (18 male and 18 female narrators) from LibriVox (a public domain collection of audiobooks) [45]. Anechoic head-related transfer functions for 6 behind-the-ear microphones (*M* = 6) in a binaural hearing aid set-up [46] (with approximately 7.5 mm distance between neighbouring microphones) were used for making directional audio. All audio was sampled at 44.1 kHz. The long term spectrum of the babble sources was matched with the average spectrum of all the target speech streams. The experiments included a condition without background babble noise (referred to as the noise-free case) and two noisy conditions with SNRs^+^ of -1.1 dB and -4.1 dB. Note that both SNRs are negative, resulting in challenging conditions where the signal power of the background babble noise is higher than the signal power of the attended speaker. Different scenarios regarding the relative position of both speakers are simulated, namely (−90° (utmost left), 90° (utmost right)), (30°, 90°), (−30°, -90°), and (−5°, 5°) with angle between speakers 180°, 60°, 60° and 10° respectively.

This RBM data set is used in all evaluation experiments throughout this paper, unless mentioned otherwise.

#### 3.1.3. DNN training dataset

A separate audio dataset was needed to train the model for the speaker independent deep learning based source separation as explained in 2.2.3. A novel dataset, called *Corpus Gesproken Nederlands Mix* (CGNM), was created. It used the *Corpus Gesproken Nederlands component-o-Flemish* (CGN-o-VL) [47], which contains 150 Dutch speakers of which 102 (51 female and 51 male) were randomly chosen to build up the training set, containing 27 hours of unique single-speaker speech in total (16 minutes per speaker on average). 20 000 2-speakers mixtures were artificially created using randomly selected single-speaker audio from CGN-o-VL. These mixtures totalled 60 hours of overlapping speech, to be used as training set. Even though in the RBM dataset (Subsection 3.1.1) only female-female mixtures are used, male-female and male-male mixtures are included in the training set as well for better generalizability. Furthermore, both speakers can be located anywhere between -100° and 100°, again to make the network more general. Two types of datasets were made: one containing the 20 000 mixtures without background babble noise (CGNM-noisefree), and the same mixtures with background babble noise (CGNM-noise). The SNR for the background noise for each mixture was uniformly sampled between -4 dB and 4 dB. Babble speakers were taken from the LibriSpeech corpus [48] and were positioned in the same way as in 3.1.1. All DNN models in this paper are trained on this CGNM dataset. Both CGNM-noisefree and CGNM-noise were used to train the DNN models, unless specified otherwise.

To assess separation quality, 12 held-out female speakers (totalling 3.2 hours of unique speech) were chosen to create a test set. The test set consisted of 728 2-speaker mixtures (totalling 2.2 hours of mixed speech). While the RBM set remains the main dataset for evaluation throughout this paper, this CGNM test set is introduced as it is more closely related to the CGNM train set, which was used to train the DNN models. Comparing DNN separation performance on the CGNM test set and the RBM set, will allow to address some generalization issues of the DNN in Subsection 4.1.

### 3.2. Data preprocessing and design choices

#### 3.2.1. EEG preprocessing

All EEG data was filtered using an equiripple bandpass filter with passband between 0.5 Hz and 10 Hz, and with passband attenuation of at most 0.5 dB and stopband attenuation of 20 dB (lower) and 15 dB (upper). Previous studies [1, 5, 49] have shown that cortical envelope tracking is best within this frequency range. EEG data was then downsampled to 40 Hz.

#### 3.2.2. M-NICA audio preprocessing

The microphone signals were lowpass filtered with an 800 Hz cut-off and downsampled to 8 kHz before computing the energy envelopes, since it was found to be beneficial for the source separation process under low SNR conditions [22]. Energy was computed every 200 samples (25 ms), bringing down the sampling rate to 40 Hz.

#### 3.2.3. Deep learning design choices

For the deep learning model utterances were also downsampled to 8 kHz. An STFT with a 32 ms window length (256 samples) and a hop size of 8 ms was used. A bidirectional long short-term memory recurrent neural network (BLSTM-RNN) with 2 layers and 600 hidden units each, was used to estimate the embeddings which where then normalized. The embedding dimension was chosen at *D* = 20 and since the number of frequency bins was Ω = 129, the total number of output nodes was *D*Ω = 20 * 129 = 2580. This is identical to the setup in [24]. For the enhancement network *g*′ another BLSTM-RNN was used with 2 layers using 300 hidden units each. The output layer had Ω = 129 nodes and a sigmoid function was applied on top to estimate the final masks. The hyperparameter *γ* in (12) is set to 1.

According to the principles of *curriculum learning* [24, 50], the networks were first trained on segments of 100 time frames, before training on the full mixture. The weights and biases were optimized with the Adam learning algorithm [51] and early stopping was used on the validation set. The log-magnitude of the STFT coefficients were used as input features and were mean and variance normalized. Zero mean Gaussian noise with standard deviation 0.2 was applied to the training features for better generalization [52]. For the single-channel scenario a single microphone was randomly selected. For the multi-channel scenario the input consisted of the average of the left-ear and the average of the right-ear microphone signals, as well as the spatial features (see (14)) from 3 microphone pairs (with one pair using a left-ear and a right-ear microphone and pair 2 and 3 using two left-ear and two right-ear microphones, respectively). All networks were trained using TensorFlow [53]. The code for the deep learning based source separation can be found here: https://github.com/JeroenZegers/Nabu-MSSS.

#### 3.2.4. MWF design choices

For the estimation of MWF coefficients an STFT with a 64 ms window length (512 samples) and a hop size of 32 ms (256 samples) was applied to the microphone signals, after they were downsampled to 8 kHz. The thresholds for estimating the VAD tracks for the different approaches were determined empirically. For energy envelopes extracted from the M-NICA and DNN-based source separation, the 25th percentile of the per sample amplitude of the envelope was used as the threshold, above which the speaker was considered to be active.

#### 3.2.5. Audio envelope extraction

The envelope extraction method from [14] was used. Each of the speech streams used as stimulus were filtered with a gammatone filterbank [54], splitting the signal into 15 filter bands. In each gammatone band, the absolute value of each sample was taken after which a power law compression with an exponent of 0.6 was applied. The power law-compressed samples were then bandpass filtered in the same manner as was done for the EEG signals resulting in subband envelopes. The subband envelopes were added together and downsampled to 40 Hz to form a single ‘powerlaw subband’ envelope. Power law compression aims to replicate the non-linear transformation of the stimulus in the human auditory system, with relatively higher attenuation of higher amplitude signals [55]. This method of auditory inspired envelope extraction has been found to result in significantly better attention decoding accuracies than other envelope extraction methods [14].

Power law subband envelopes were only used in cases where broadband speech streams where available, i.e., the DNN and MWF envelopes built from the speech streams obtained with the DNN-based mask (15) or the MWF-based mask (7), respectively. Since M-NICA directly extracts energy envelopes from the broadband speech streams, the gammatone filterbank could not be applied here. When applying AAD on the M-NICA-envelopes, the square-root of the extracted energy envelopes was used to transform them to an amplitude envelope, since energy envelopes are sub-optimal to perform AAD [14].

#### 3.2.6. AAD design choices

In order to perform AAD, EEG data was split into trials of 30 s, resulting in 276 trials per subject for each of the 3 different angles between speakers (separations of 180° for (−90°,90°), 60° for (30°,90°) and (−30°,-90°), and 10° for (−5°,5°) respectively). Decoders were trained (using leave-one-trial-out (LOO) cross validation) to reconstruct the envelope of the attended speech stream. To improve decoding accuracy and eliminate the need for regularization, a single decoder was computed over the entire training set data as in [14], instead of averaging over per-trial decoders in the training set as in [2].

## 4. Results

### 4.1. Single-channel and multi-channel source separation

In some cases, for hardware reasons, a single-channel set-up can be preferred over a multi-channel set-up. The deep learning approaches to noise-free source separation, both for single-channel and multi-channel setup, were trained on the CGNM-noisefree train set and evaluated on the CGNM-noisefree test set and the RBM noise-free set. Results are expressed in Signal-to-Distortion Ratio (SDR) [56] improvements and are shown in table 1.

**Table 1:** SDR improvement results for a single-channel and multi-channel DNN without noise.

|  | noise-free CGNM test | noise-free RBM |
| --- | --- | --- |
| single-channel | 10.0 dB | 3.8 dB |
| multi-channel | 12.3 dB | 12.3 dB |

Notice that for the single-channel case, performance on the noise-free RBM dataset is much worse than on the CGNM-noisefree test set, even though both consist of held-out female speakers. The DNN generalizes poorly to other datasets. A possible reason for this could be differences in the recording set-up, and the fact that the CGNM-noisefree test set consists of volunteers, while the RBM dataset consists of children stories told by professional story-tellers. It is concluded that the single-channel setup does not generalize well across data sets. Multi-channel separation quality is considerably better than the single-channel counter part. Furthermore, the SDR results on the CGNM-noisefree test set and the RBM dataset are also more alike, compared to the single-channel results. The network uses spatial information to characterize the speakers and does not need to solely rely on the difference in speaker characteristics to separate the speakers, as is necessary in the single-channel case [58]. In table 2, the SDR results on the RBM dataset are split up per angle between speakers. For the single-channel setting, it is observed that the model performs much better for the 180° case compared to the other cases. For the 180° case the speech amplitudes of both speakers will differ more due to head shadow effects, which may explain why it is easier for the network to separate both. These head shadow effect might also happen, although less pronounced, for the 10° (speakers at -5° and 5°) case, which could explain why it performs slightly better than the 60° (speakers at 30° and 90°) case, where both speakers are on the same side of the head. Further research is needed to support these claims/assumptions.

**Table 2:** SDR improvement results for a single-channel and multi-channel DNN for the noise-free RBM set, depending on the difference in speaker positions.

|  | 10° | 60° | 180° | avg. |
| --- | --- | --- | --- | --- |
| single-channel | 2.5 dB | 1.7 dB | 7.3 dB | 3.8 dB |
| multi-channel | 12.7 dB | 11.6 dB | 12.5 dB | 12.3 dB |

The absolute envelopes, extracted from the outputs of both single-channel and multi-channel DNNs performing speaker separation of the stimuli in the RBM dataset for the noise-free condition, were used for auditory attention decoding of the EEG data collected under matching acoustic conditions and stimuli. Figure 2 shows AAD accuracies over 18 subjects under the noise-free condition, for the 3 different angles between the speakers. For all angles between speakers, the multi-channel DNN resulted in significantly higher AAD accuracies compared to the single-channel DNN, as was expected from the results in table 1. We therefore continue with the multi-channel approach for all further analyses.

**Figure 2:**
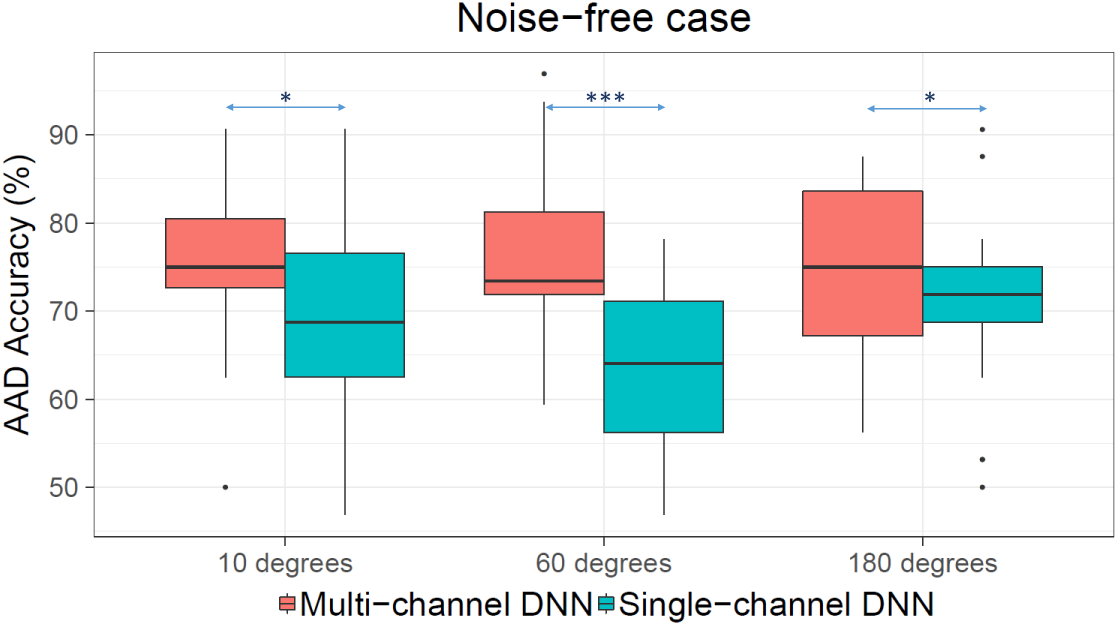
AAD accuracies (on 30 s trials) of 18 subjects listening in noise-free conditions when envelopes from the output of multi-channel DNN and single-channel DNN were used. Comparisons between the different approaches were done using Wilcoxon’s signed-rank test [57] with Holm correction. : ‘***’ for p < 0.001, ‘*’ for p < 0.05.

### 4.2. Linear versus non-linear source separation for AAD

The per-subject AAD accuracies on the RBM dataset were computed for each of the 3 different angles between speakers and the 3 noise conditions. For each 30 s trial, the correlation of the reconstructed attended envelope was computed with 5 pairs of envelopes extracted using different speech enhancement methods:

- ‘Oracle’ - powerlaw subband envelopes extracted from the original speech signals (not mixed)
- ‘M-NICA’ - amplitude envelopes extracted from the output of the M-NICA source separation algorithm (red dashed track in Figure 1)
- ‘DNN’ - powerlaw subband envelopes extracted from the output of the DNN-based source separation algorithm (also red dashed track in Figure 1).
- ‘M-NICA+MWF’ - powerlaw subband envelopes extracted from the output of MWFs when using the output of the M-NICA based source separation algorithm to build VAD tracks (green track in Figure 1).
- ‘DNN+MWF’ - powerlaw subband envelopes extracted from the output of MWFs when using the output of the DNN based source separation algorithm to build VAD tracks (also green track in Figure 1).

Figure 3 shows the resulting AAD accuracies for the 18 subjects for the noise-free condition, for -1.1 dB and for -4.1 dB. For each angle between speakers, comparison between the different approaches were done using Wilcoxon’s signed-rank test with Holm correction. The green boxplots correspond to the green lines (MWF envelopes), and the red box plots correspond to the dashed red lines (M-NICA/DNN envelopes) respectively in Figure 1.

**Figure 3:**
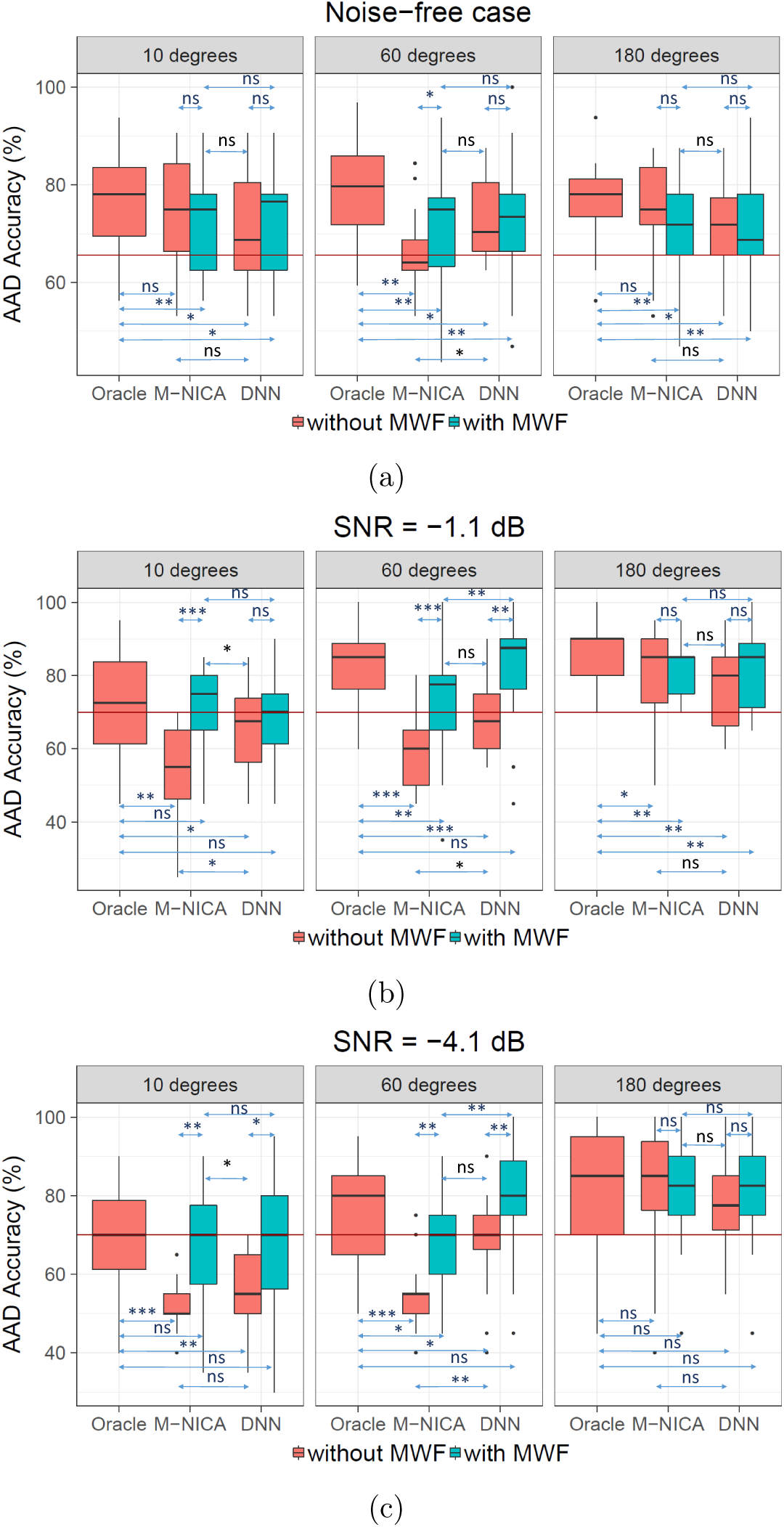
AAD accuracies (on 30 s trials) of 18 subjects for different speech separation approaches for 3 cases: the noise-free case, -1.1 dB SNR, and -4.1 dB SNR. The results of using the M-NICA and DNN approaches are compared with the results when using oracle speech envelopes for AAD. The separated streams were either directly used for AAD (red boxes here and dashed red lines in Figure 1), or used to generate VAD tracks for MWFs (green boxes here and green lines in Figure 1), the outputs of which were then used for AAD. The red line indicates the significance level. Comparison between the different approaches were done using Wilcoxon’s signed-rank test with Holm correction. : ‘***’ for p < 0.001, ‘**’ for p < 0.01, ‘*’ for p < 0.05, ‘ns’ for no significant difference.

An interesting observation from Figure 3 is that a training-free linear algorithm as proposed in [22] (here represented by ‘M-NICA+MWF’) is able to perform at least on par and often even better than a pre-trained complex deep model (here represented by ‘DNN’) in all speaker angle and noise conditions. When breaking down the subcomponents of [22] into M-NICA and MWF separately, or when using the DNN as support for the MWF, some other interesting observations can be made, which are briefly reported in the remainder of this subsection.

In the noise-free condition, for 180° and 10° separations, the M-NICA envelopes result in accuracies that match those when using oracle speech envelopes for AAD while the DNN envelopes perform significantly poorer. On the other hand, for 60° speaker separation, the DNN envelopes outperform M-NICA envelopes, yet upon including the MWF filtering step, there are no more significant differences between the 2 source separation approaches. However, the results are still significantly lower than with oracle speech envelopes. It must be noted that in the 60° case, the competing speaker positions are on the same side of the listener’s head ((−30°,-90°) or (30°,90°)). It has already been observed in [22] that the same 60° case was challenging for M-NICA, indicating difficulty to extract discriminative spatial features under such conditions

Figure 3 also shows AAD accuracies from trials in a noisy condition with -1.1 dB SNR. For 180° speaker separation, it can be seen that all source separation approaches, with or without the MWF filtering step, result in similar AAD performances, but do not match up to that of oracle speech envelopes. Nevertheless, the performance decrease is minimal, in particular when an MWF is used. For 10° and 60° speaker separation, the DNN envelopes result in better performance than the M-NICA envelopes. MWF filtering substantially improves results for both approaches for the 60° case, where the DNN+MWF combination even matches performance with the oracle speech envelopes. For the 10° case, only the M-NICA approach requires MWF filtering to match oracle speech performance.

For the -4.1 dB SNR (Figure 3), which is a highly challenging condition, it is observed that for 180° separation, the performances of all the approaches do not differ significantly from each other. However, for the 10° condition, the M-NICA and DNN envelopes result in accuracies below chance level, and only with the additional MWF filtering, do the results match up to those of oracle speech envelopes. In the 60° case, the MWF filtering does improve results for both the source separation approaches, however only the DNN+MWF approach’s results match up to those of oracle speech envelopes.

### 4.3. Speech enhancement performance

To estimate the effectiveness of the MWF for speech enhancement, we looked at the improvement in the signal to interference-plus-noise ratios (SINRs) and PESQ, on the RBM dataset, at the output of the MWFs when using VAD tracks generated from 3 sources: the oracle speech envelopes, the M-NICA envelopes, and the DNN envelopes. The SINR is defined as the ratio of the power of the attended speaker to the total of the power of the unattended speaker and the background noise. The reference input SINR is taken as the highest SINR among the 6 microphones (note that the achieved SINR improvement will be larger for the other microphones). For each noise condition and angle between speakers, the improvement in SINR was computed for 8 story parts to which the subjects had to listen without interruption. Figure 4 shows the improvement in SINRs of the 8 story parts, for the 3 noise conditions and 3 different angles between speakers. For the noise-free case, it is observed that the oracle speech envelopes result in more speech enhancement than M-NICA or DNN envelopes in all separation angles (although perceptually there is not much difference: the difference between 40dB and 20dB SINR is hard to notice). However, in the presence of background noise, the DNN+MWF approach significantly outperforms the M-NICA+MWF approach, and matches up to the performance with the oracle speech envelopes in the 10° and 180° cases. For the acoustically more challenging case of 60° (since the speakers are on the same side of the head), it is observed that the DNN+MWF approach performs almost similar to the oracle+MWF approach (although the small differences are still statistically significant), and both of them substantially outperform the M-NICA+MWF approach.

**Figure 4:**
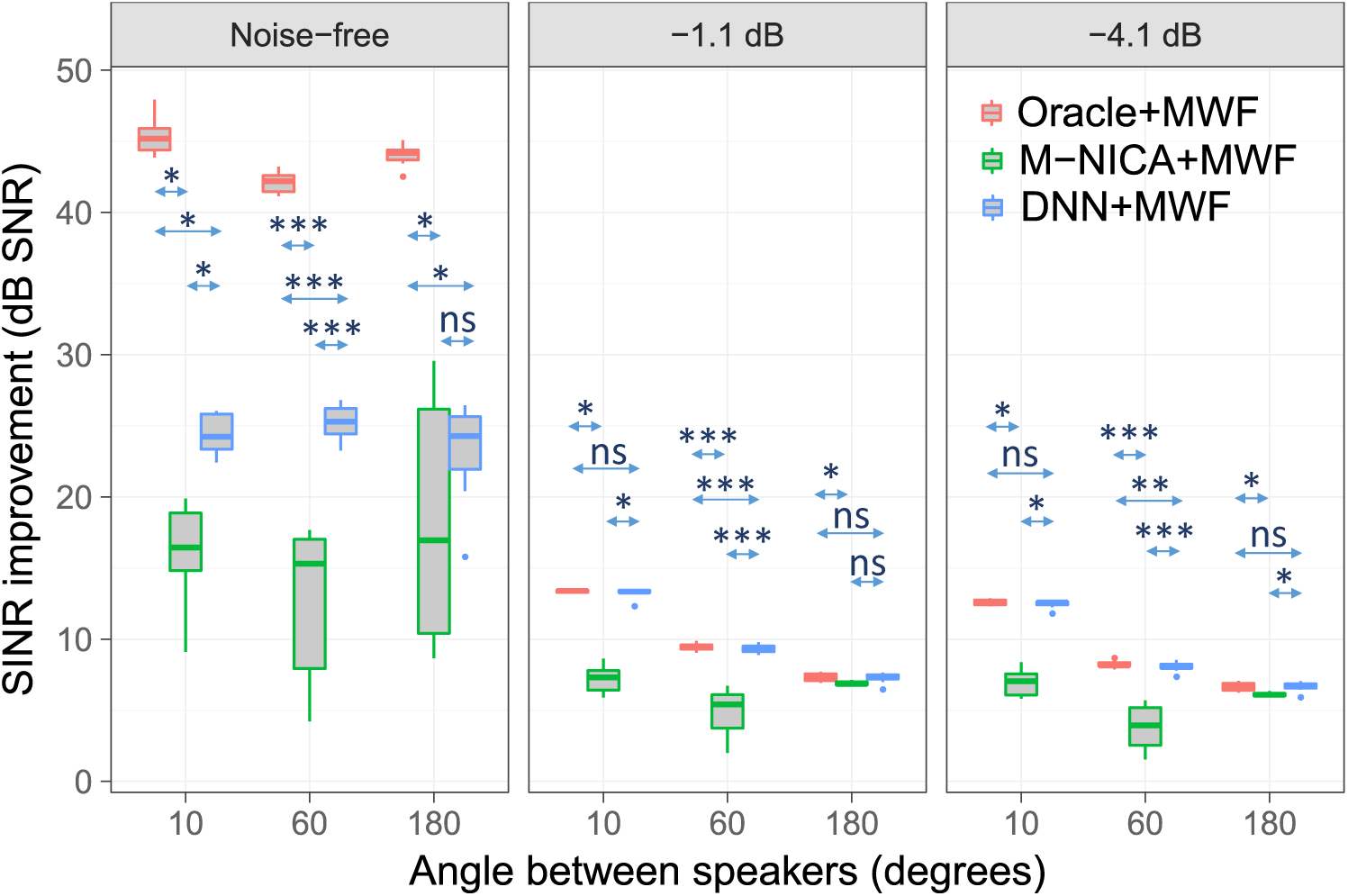
Boxplots showing improvement in signal to interference-plus-noise ratios of 8 story parts when the MWFs use VAD tracks extracted from the different speech separation approaches. The results of using M-NICA and DNN outputs for the VAD are compared with the results when using oracle speech for the VAD. Comparison between the different approaches were done using Wilcoxon’s signed-rank test with Holm correction. : ‘***’ for p < 0.001, ‘**’ for p < 0.01, ‘*’ for p < 0.05, ‘ns’ for no significant difference.

The results for PESQ, a measure that focuses more on speech intelligibility, are shown in Figure 5. The significance tests are very similar to those of SINR and thus the above observations for SINR also hold for PESQ.

**Figure 5:**
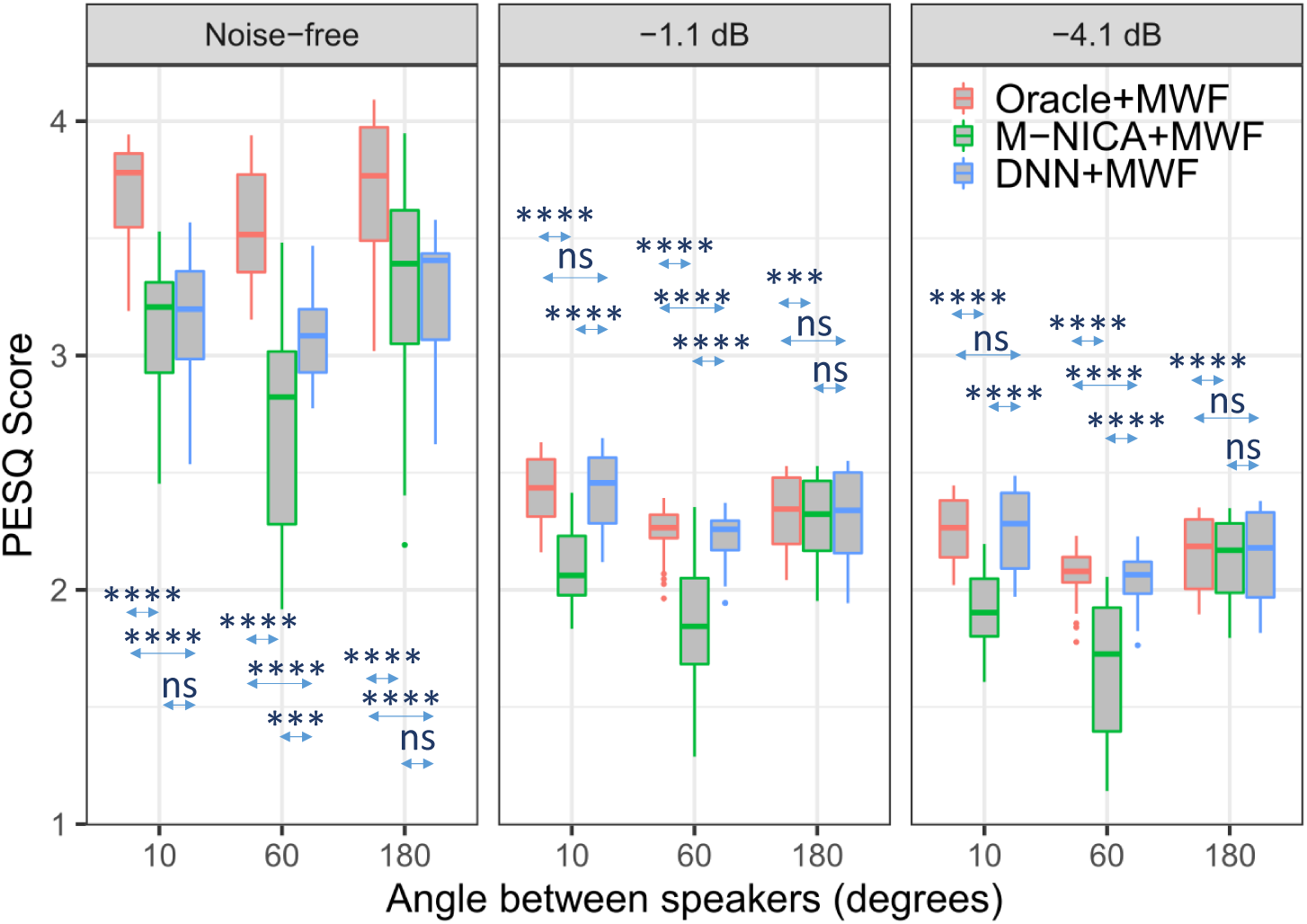
Boxplots showing PESQ scores of 8 story parts when the MWFs use VAD tracks extracted from the different speech separation approaches. The results of using M-NICA and DNN outputs for the VAD are compared with the results when using oracle speech for the VAD. Comparison between the different approaches were done using Wilcoxon’s signed-rank test with Holm correction. : ‘****’ for p < 0.0001, ‘***’ for p < 0.001, ‘**’ for p < 0.01, ‘*’ for p < 0.05, ‘ns’ for no significant difference.

### 4.4. Gain control performance

AAD accuracies are dependent on evaluated trial lengths, which was set to 30 s in our experiments. In the context of gain control in a neuro-steered hearing device, longer trial lengths can result in higher accuracies, but at the cost of slow decisions affecting the speed with which the device can follow a switch in attention of the user. The minimal expected switching duration (MESD) [59] is a performance metric for AAD algorithms that takes this trade-off into account. This metric finds the optimal operating point to minimize the expected time required for such a hearing device to switch between speakers in a robust fashion after the user switches attention. The MESD toolbox [60] was used to compute the MESDs of the best performing algorithm (DNN+MWF) and the oracle case for the 3 different angles between speakers and the 3 noise conditions. The results are shown in Figure 6. Using Wilcoxon’s signed-rank test with Holm correction, we found that the MESDs of the oracle case were not significantly lower than those of DNN+MWF, except for all speaker separations in the noise-free condition (*p* = 0.05), and for 180° at -1.1 dB SNR (*p* < 0.05). For all other cases, there were no significant differences between the two approaches.

**Figure 6:**
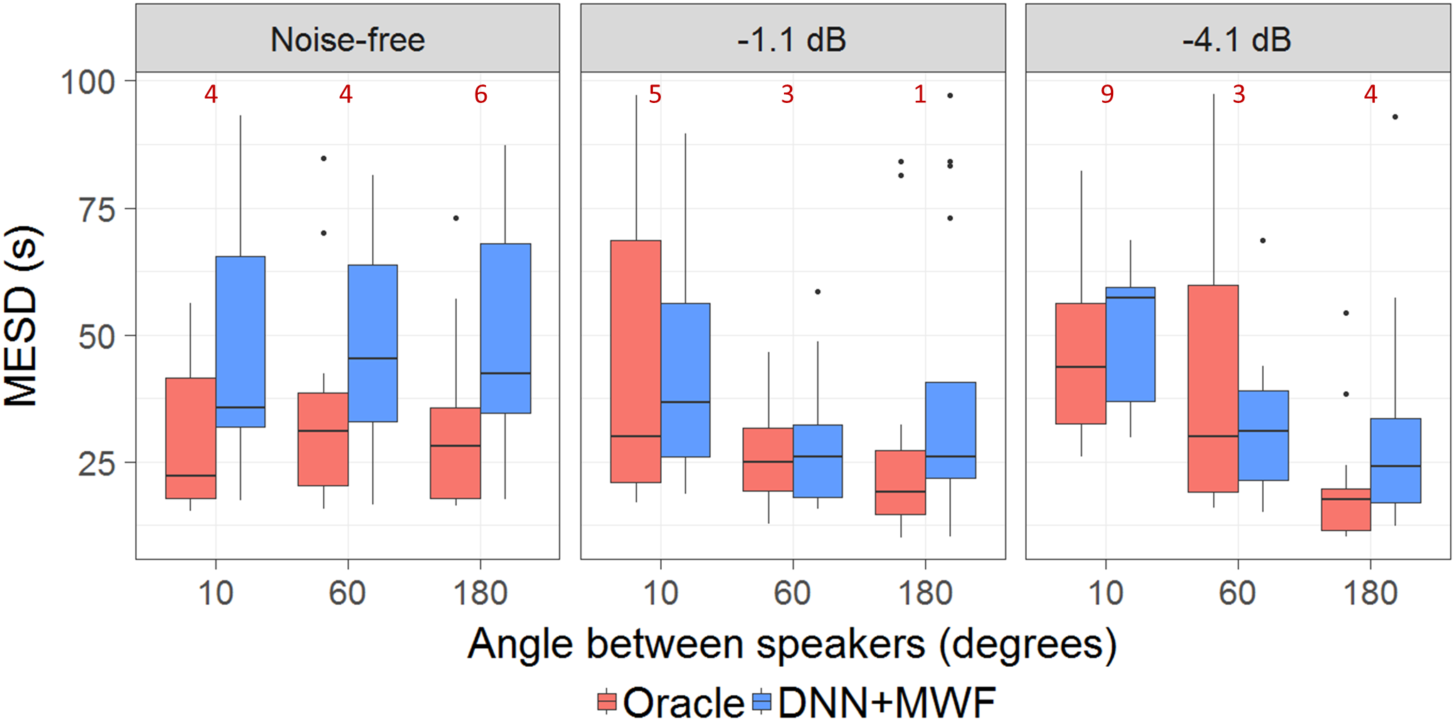
Boxplots showing MESDs for two cases: when attention decoding was performed using ‘Oracle’ speech envelopes versus when using ‘DNN+MWF’ speech envelopes. The numbers in red indicate the number of outliers (MESD > 100s) which were removed from the plot (such high values are due to near-chance level performances, which are too low to achieve an acceptable gain control, leading to extremely high MESDs). They were included in the statistical analysis.

## 5. Discussion

Previous studies that combine speech enhancement/separation with AAD investigated only one type of algorithm (either linear [20, 22, 23] or DNN-based [31, 32]), mostly in relatively mild acoustic conditions. Furthermore, each study has its own specific set-up (single-vs. multi-microphone, ECoG vs. EEG, matched or mismatched acoustic conditions during EEG recordings, etc.) which does not allow to directly compare the results achieved in these studies. The study in this paper was designed to investigate the feasibility of a neuro-steered speech enhancement pipeline in challenging acoustic conditions with negative SNRs and various speaker positions, while also comparing different source separation approaches to produce the speech envelopes used by the attention decoding block. We investigated the AAD performance of a training-free and purely linear audio signal processing algorithm (M-NICA+MWF) on the one hand, and a non-linear DNN on the other hand, as well as a combination of a linear with a non-linear approach (DNN+MWF), where the latter acts as a supporting voice activity detector for the former. In the remainder of this section, we will discuss the main findings of our study.

### 5.1. Multi-microphone outperforms single-microphone

Existing AAD studies using DNNs for the acoustic source separation were based on single-microphone recordings [31, 32]. We observed that a single-channel approach generalizes less well across datasets and that a multi-channel approach is more robust in terms of separation quality. In addition, envelopes extracted from the output of the multi-channel neural network resulted in AAD accuracies that were more robust to varying speaker positions compared to those from the single-channel neural network. It is concluded that a multi-microphone set-up is crucial to obtain a sufficiently high AAD accuracy in practical settings. Thus, we chose the multi-channel neural network approach for our comprehensive analysis of the neuro-steered source separation.

### 5.2. Linear algorithm outperforms DNN

In the literature, several algorithms have been proposed to combine AAD with speech separation methods. Concerning the speech separation part, two main strategies can be distinguished; those based on traditional (linear) beamforming approaches [20, 22, 23] and those based on (non-linear) neural networks [31, 32]. However, these two strategies have never been compared to each other in terms of the resulting AAD performance. The results in Section 4 demonstrate that the AAD performance with a linear speech separation algorithm (represented by ‘M-NICA+MWF’ in figure 3) is at least on par and often outperforms a non-linear DNN approach (represented by ‘DNN’ in figure 3) in all investigated conditions. This holds both in terms of AAD performance (Fig. 3) and speech enhancement performance (Fig. 4). In addition, the linear method is computationally cheaper than the DNN approach (in terms of operations per second and memory size), while also being training-free thereby not depending on representative training data. Furthermore, implementing the MWF as an adaptive, causal, low-latency filter is relatively straightforward, and latencies can be limited to less than 5 ms [61] (see also Section 5.4 below). Speech separation based on deep clustering on the other hand, usually uses a non-causal network where the computation of the embedding vector at present time is based on all past and future time samples, although recently also causal DNNs for speech enhancement have been proposed [32, 62].

### 5.3. Combining the best of both worlds: DNN and MWF

While linear and non-linear approaches both individually result in good speech separation, a combination of both - as in the DNN+MWF approach - resulted in the best AAD performance, particularly in challenging acoustic conditions. In this case the (non-linear) DNN is used as a VAD mechanism to inform the (linear) MWF which performs the actual speaker separation and denoising, thereby combining the best of both worlds. This results in the best AAD performance as well as the highest SINR improvement.

### 5.4. Real-time aspects

When using a weighted overlap-add (WOLA) procedure [63], the algorithmic delay of the MWF is equal to the window length used in the STFT computations, i.e., the number of samples over which the discrete Fourier transform is computed. For example, for a hearing aid using a sampling rate of 20480 Hz and a 96-point STFT windowing [61], the algorithmic delay is less than 5 ms. Note that a delay in the VAD information (e.g. due to possible non-causality of the BSS block in Figure 1), does not have an impact on the overall input-output delay of the stimulus presented to the user. This is because the adaptation of the MWF filter coefficients can be decoupled from the actual filtering operation itself, where the former can lag behind on the latter. This implies that any delay or non-causality in the M-NICA or DNN block will not create a bottleneck towards real-time speech processing. It will only add some inertia in the updating of the MWF coefficients to adapt to changes in the acoustic scenario.

### 5.5. EEG-based AAD is feasible in challenging low-SNR conditions

The SNRs in this study were chosen such that the speech intelligibility varied across conditions. The estimated speech recognition threshold (SRT), at which a normal-hearing subject can understand 50% of the attended story on average, was found to be -7.1 dB (see [43] for details of the estimation procedure). The SNRs -4.1 dB and -1.1 dB were chosen by adding 3 dB steps from the 50% speech intelligibility point. The -7.1 dB condition was excluded from this study due to the difficulty subjects faced in focusing on the attended speaker (as reflected in the AAD accuracies under this condition reported in [43]). Subjective speech intelligibility reported during the recording sessions for the different angles between speakers show that -4.1 dB and -1.1 dB SNRs also are not easy listening conditions. Compared to [23] where the noise conditions are relatively mild (4 dB and 9 dB SNR), [32] where there is no background noise, and [22] where the EEG recording conditions did not match the acoustic conditions, our current study provides a holistic view on the performance of a neuro-steered speech enhancement pipeline under realistic to challenging acoustic conditions. It is found that not only EEG-based AAD with no access to clean-speech sources is feasible under challenging acoustic conditions, but also that the AAD accuracies and improvement in SINRs for DNN+MWF get remarkably close to oracle performance.

### 5.6. Future outlook

The proposed neuro-steered speech enhancement and its analysis over a range of acoustic conditions is a step towards effectively incorporating attention decoding and source separation algorithms in neuro-steered hearing prostheses. With the field of deep learning expanding and exerting its presence in various scientific domains, we have tried to address the possibility of deep learning based source separation in this context and systematically compare it with a classical signal processing approach. While the realisation of neuro-steered hearing prostheses is still far from being a reality, there are ongoing advancements that can accelerate this process. In addition to hardware requirements such as miniaturisation of neural recording equipment, better computational and storage capabilities in hearing devices etc., also newer approaches for improved attention decoding itself based on DNNs [11–13], canonical correlation analysis [64] or state-space modelling [3, 10] can contribute to faster and more robust attention decoding and hence better neuro-steered speech enhancement. Finally, a limitation of this study is that it does not include reverberant conditions, which can both affect speech separation algorithms and the AAD performance.

## 6. Conclusion

In a multi-speaker ‘cocktail party’ scenario, a hearing aid’s noise reduction algorithm does not have the knowledge of which speaker the user intends to listen to. Incorporating AAD algorithms to detect the attention of the user, leads to the concept of so-called neuro-steered hearing aids. In this study a neuro-steered speech enhancement pipeline is presented, without access to clean speech signals in challenging acoustic scenarios.

The performance of a linear as well as DNN-based speaker separation approach was analyzed. It was found that a purely linear approach often outperforms the DNN-based approach. However, the best performance, under challenging scenarios, was obtained when combining the best of both worlds, i.e., when using the DNN to provide VAD information to a linear data-driven beamformer. In addition, for the DNN-based speaker separation, the benefit of using a multi-microphone system compared to a single-microphone system was demonstrated. An additional advantage of using a linear data-driven beamformer is that it decouples the algorithmic delay in the blind source separation of the audio signals from the total system delay, thus facilitating robust real-time speech processing. With this study, we present a proof-of-concept of feasibility of a neuro-steered speech enhancement pipeline in challenging acoustic conditions.

## Acknowledgments

The work is funded by FWO project nr G0A4918N, the Flemish Goverment (AI Research Program) and the ERC (637424 and 802895) under the European Union’s Horizon 2020 research and innovation programme. Jeroen Zegers is supported by an SB PhD scholarship 1S66217N of FWO. The scientific responsibility is assumed by its authors.

## Footnotes

‡ Note that both AAD and speaker separation are expected to be more difficult when the competing speakers are of the same gender.

§ For the sake of conciseness, the frequency variable *ω* is omitted for autocorrelation matrices in this section.

‖ There is also the possibility of using other parameters, speech presence probability [38] for instance, to make softer VADs, but this is not within the scope of this paper.

¶ This corresponds to applying the mask to the output of a forward-steering beamformer. Other choices are possible (e.g. select only one microphone, or summing magnitude spectra and adding the phase of one of the microphones), but these were empirically found to not have a significant impact on the results.

+ In this paper (input) SNR is defined as the ratio of the power of the target speaker to the power of background babble. Note that the actual SNR is even lower due to the presence of an interfering speaker, which is not included in this SNR metric.

